# Model Parameter identification using 2D vs 3D experimental data: a comparative analysis

**DOI:** 10.1101/2023.05.17.541071

**Authors:** Marilisa Cortesi, Dongli Liu, Christine Yee, Deborah J. Marsh, Caroline E. Ford

## Abstract

Computational models are becoming an increasingly valuable tool in biomedical research. They enable the quantification of variables difficult to measure experimentally, an increase in the spatio-temporal resolution of the experiments and the testing of hypotheses.

Parameter estimation from *in-vitro* data, remains a challenge, due to the limited availability of experimental datasets acquired in directly comparable conditions. While the use of computational models to supplement laboratory results contributes to this issue, a more extensive analysis of the effect of incomplete or inaccurate data on the parameter optimization process and its results is warranted. To this end, we compared the results obtained from the same *in-silico* model of ovarian cancer cell growth and metastasis, calibrated with datasets acquired from two different experimental settings: a traditional 2D monolayer, and 3D cell culture models.

The differential behaviour of these models will inform the role and importance of experimental data in the calibration of computational models’ calibration. This work will also provide a set of general guidelines for the comparative testing and selection of experimental models and protocols to be used for parameter optimization in computational models

**Author summary:** Parameter identification is a key step in the development of a computational model, that is used to establish a connection between the simulated and experimental results and verify the accuracy of the *in-silico* framework.

The selection of the *in-vitro* data to be used in this phase is fundamental, but little attention has been paid to the role of the experimental model in this process. To bridge this gap we present a comparative analysis of the same computational model calibrated using experimental data acquired from cells cultured (i) in 2D monolayers, (ii) in 3D culture models and (iii) a combination of the two.

Data acquired in different experimental settings induce changes in the optimal parameter sets and the corresponding computational model’s behaviour. This translates in a varying degree of accuracy during the validation procedure, when the simulated data are compared to experimental measurements not used during the calibration step.

Overall, our work provides a workflow and a set of guidelines to select the most appropriate experimental setting for the calibration and validation of computational models.

## Introduction

Computational models (CMs) are becoming an increasingly important tool in biomedical research, allowing for the study of complex phenomena in controlled environments [1, 2], the prediction of a system’s behaviour in multiple conditions [3, 4], and the testing of hypotheses [5, 6]. Experimental corroboration, here defined as the combination of CM calibration and validation, is a key aspect in the development of these tools, as it represents the connection between the *in-silico* and the *in-vitro* models.

Calibrating a CM consists of the identification of its parameters so as to recapitulate the process of interest. Multiple search and optimisation algorithms can be used in this phase [3, 7–9], although empirical parameters selection remains common when a small number of well constrained parameters needs to be identified. CM validation is the procedure used to determine the simulations’ information content through a quantification of each simulation’s accuracy. This step is widely recognised as fundamental for the development of useful and effective CMs and a wealth of resources and guidelines are available from the recent scientific literature [4, 10, 11].

The effect of the experimental model (EM) on the results of the model’s corroboration, however, remain largely unexplored. Difficulties in acquiring or accessing datasets containing all the relevant information have been a major obstacle, especially when relying solely on the literature. While this issue is becoming less relevant, due to an increase in open access data availability and more extensive collaboration between wet-lab and dry-lab researchers, the widespread use of 3D cell culture models is an aspect that warrants further investigation. Indeed, changing the setup has been shown extensively to affect cell behaviour and results of *in-vitro* experiments [12–14]. As such, the corroboration of a single CM using data acquired on multiple EMs has potentially deleterious effects on the accuracy and reliability of the simulated results. To test this hypothesis, we here present a comparative study of the same CM corroborated with datasets acquired using either a 2D monolayer culture, 3D EMs, or a combination of the two.

As a case study, we chose to focus on transcoelomic metastasis, the major mechanism of metastasis (or cancer spread) in ovarian cancer [15]. It occurs via the seeding of cancer cells onto the omentum, or other tissues within the abdominal cavity, following their detachment from the ovary and it is enabled by the ascites fluid which builds up in this region, and by the receptiveness of the surrounding tissues to colonisation [16–22].

The extensive cell-cell and cell-environment interactions involved in this process have led to the development of a number of 3D cell culture models in order to research facets of this phenomenon [12, 23–27]. We selected a 3D organotypic model [28, 29] used extensively to study the invasion and adhesion capabilities of ovarian cancer cells [30–34] and 3D bio-printed multi-spheroids for the quantification of proliferation.

Together, these EMs allow us to study both the initial phases of metastasis, when cells floating within the abdominal cavity exhibit a phenotype associated with very little proliferation, and the later stages of this process, when sustained cancer cell proliferation is observed within the omentum [35]. In all cases, standard assays performed on 2D monolayers were used as comparison.

## Materials and methods

### Cell Culture

The high-grade serous ovarian cancer (HGSOC) cell line PEO4 was used for this study [36]. This cell line is characterised by resistance to platinum treatment and can be considered a good model of recurrent disease [37]. Cells were kindly gifted by Dr Simon Langdon (University of Edinburgh, Edinburgh, UK) and labelled with GFP (pLKO.1-Neo-CMV-tGFP vector from Sigma-Aldrich, USA) to enable their identification within the 3D organotypic model. Cells were maintained in RPMI medium (Thermo Fisher, Waltham, MA, USA), supplemented with 10% FBS (Sigma-Aldrich, USA), 1% Pen-strep (Sigma-Aldrich, USA) and 1% GlutaMAX (Thermo Fisher, Waltham, MA, USA).

The 3D organotypic model, chosen to evaluate adhesion and invasion, was built co-culturing PEO4 cells with healthy omentum-derived fibroblasts and mesothelial cells collected from patients undergoing surgery for benign or non-metastatic conditions patients at the Royal Hospital for Women and Prince of Wales Private Hospital (site specific approval ethics # LNR/16/POWH/236). The South Eastern Sydney Local Health District Human Research Ethics Committee (SESLHD HREC approval #16/108) approved the collection of these samples. The protocol for the realization of the organotypic model is fully described in [38]. In brief, 100 µl of a solution of media, fibroblast cells (4 ·10^4^ cells/ml) and collagen I (5 ng/µl, Sigma-Aldrich, USA) was added to the wells of a 96-well plate. After 4 hours of incubation at 37°C and 5% CO_2_, 50 *µ*l of media containing 20,000 mesothelial cells was added on top. The whole structure was maintained in standard culturing conditions for 24 h prior to seeding of cancer cells. PEO4 cells were added at a density of 1·10^6^ cells/ml (100 *µ*l/well) in 2% FBS media.

Proliferation was quantified in 3D multi-spheroids encapsulated in PEG-based hydrogels created using the Rastrum 3D bioprinter (Inventia Life Science, Alexandria, New South Wales) [39, 40]. Three-thousands PEO4 cells per well were printed as an “Imaging model” using the Px02.31P matrix, atop an inert hydrogel base, and across an entire tissue culture-grade flat bottomed 96-well plate. The hydrogel matrix is characterised by a 1.1 kPa stiffness and by its functionalisation with arginylglycylaspartic acid (RGD), a peptide shown to promote cell adhesion [41]. Printed spheroids were maintained at 37°C and 5% CO_2_ for a week prior to each experiment.

### Proliferation

Proliferation in 2D was measured via MTT assay (Thermo Fisher, Waltham, MA, USA), following the manufacturer’s protocol. Briefly, PEO4 cells were seeded in 96 wells plates at a density of 10,000 cells per well. After 24 h, treatment with different concentrations of either cisplatin (50, 25, 12.5, 6.2, 3.1, 1.6, 0.8, 0.4, 0 *µ*M) or paclitaxel (50, 25, 12.5, 6.2, 3.1, 1.6, 0.8, 0.4, 0 nM) was administered. Following 72 h of treatment a solution of 2 mg/ml of MTT was added to each well and incubated for 3 hours. The media-MTT solution was then discarded, and the formazan crystals solubilized in DMSO (Sigma-Aldrich, USA). Absorbance was measured at 570 nm. All data were normalised with respect to the untreated condition and corrected for the absorbance of RPMI medium. A total of 3 biological replicates, each comprising 3 technical replicates was analysed for condition.

Real-time monitoring of PEO4 cell growth within the hydrogel multispheroids and in the absence of treatment was conducted using an IncuCyte S3 Live Cell Analysis System (Sartorius, Gottingen, Germany). Three cell densities (2000, 3000 and 4000 cells/well in hydrogel) were considered, and the phase count function of the device’s analysis software was used to determine the number of cells, every hour over 7 days. A total of 10 wells/condition were considered for this experiment. An evaluation of confluency with CellTiter-Glo 3D (Promega, Madison WI, USA) was also conducted at the end of monitoring. Real-time monitoring and the end-point assays were in agreement and a density of 3000 cells per well was chosen for further experiments with only CellTiter-Glo 3D.

Treatment with cisplatin and paclitaxel, in the same concentrations used for the 2D experiments, was administered 7 days after the printing, to allow for the establishment of a stable 3D culture. Measurements were conducted after 72 h following the manufacturer’s protocol. A total of 3 biological replicates, each comprising at least 3 technical replicates, was analysed for each condition. All data were corrected with respect to the signal produced by the matrix devoid of cells and normalised with respect to the average value measured for the untreated control.

### Adhesion

Adhesion in 2D was evaluated as in [42]. Briefly the wells of a 96-well plate were coated with 10 µg/ml of collagen I (Sigma-Aldrich, USA) or 3% BSA (Sigma-Aldrich, USA). 100,000 PEO4 cells were seeded on top of each coating and incubated for 2, 3 or 4 h at 37°C and 5% CO_2_. Unattached cells were then washed away, prior to fixing with 96% ethanol and staining with 1% crystal violet. Cells were then lysed with 50% acetic acid and their density was quantified with an absorbance measurement (at 595 nm). A total of 3 biological replicates each comprising 2 technical replicates was considered for this analysis. Adhesion in 3D was quantified following the same procedure, just substituting the collagen coating with the organotypic model [38]. A limitation of this approach is that the measured absorbance integrates the signal from all the cell types present in the organotypic model. The amount of mesothelial and fibroblast cells is however assumed to be constant throughout the experiment.

### Invasion

Invasion in 2D was measured using Matrigel-precoated transwell chambers (Corning Life Sciences, USA). PEO4 cells were harvested with trypsin and diluted to a concentration of 1·10^6^ cells/ml in 1% FBS media. 100 *µ*l of the cell solution was added to each transwell insert and incubated for 48 h.

Following the incubation, transwell inserts were gently washed with PBS and fixed in 4% paraformaldehyde for 10 minutes. Wells were washed again in PBS and mounted on a microscope slide using DAPI mounting medium (Fluoroshield, Sigma-Aldrich, USA). Slides were let dry for at least 1 h prior to imaging at the microscope (Leica DM 2000 LED fitted with a Leica DFC450c camera). Ten images from different regions of the slide were acquired and the number of invaded cells counted using custom-made software further described in the next section. A total of 3 biological replicates each comprising 2 technical replicates was considered for this analysis.

Cancer cell invasion in 3D was measured in a substantially equivalent way, by substituting the Matrigel-precoated chambers with regular transwell inserts (Corning Life Sciences, USA) in which the organotypic model had been seeded [38]. Specific modifications in the counting software used for the 2D analysis allowed for the identification of the invaded cancer cells, which were producing a stable GFP signal.

### Invasion quantification software

The number of cells in each image was quantified through custom-made software written in Python (v.3.9) and freely available at https://github.com/MarilisaCortesi/cell counter. It uses the Otsu’s method to segment the nuclei and a labelling routine to identify each segmented region and thus determine the total number of cells (S1 Fig a., b.).The accuracy of this method was evaluated by comparing the number of cells retrieved by the software for each image, with the corresponding manual count obtained by an expert user using ImageJ (S1 Fig c.). The two measures are highly correlated (R^2^ = 0.95), and their average percentage error is consistent with the inter-operator variability for this assay (about 18%, [43]). As such, the automatic count was considered to be equivalent to the manual one.

For the images obtained during 3D experiments, two additional filters based on the average fluorescence intensity and area were used to separate cancer cells from fibroblasts and mesothelial cells. In particular, a cell was labelled as PEO4 if its area was between 50 and 5,000 pixels and its average fluorescence intensity was higher than that of the background (S1 Fig d., e., f.).

### SALSA modeling and computational simulations

Computational simulations were conducted in SALSA [44–46]), a hybrid continuous-discrete cellular automaton freely available at https://www.mcbeng.it/en/category/software.html. A cubic 3D lattice constitutes the main structure of the simulator (Fig 1a). Cells can be positioned at each of the grid’s nodes and the values of the continuous variables are computed at the same locations. Specifically It relies on specifically formatted configuration files (available as supplementary material S1 File) are used to formalize cell behaviour and initialise the experimental conditions.

**Fig 1.**
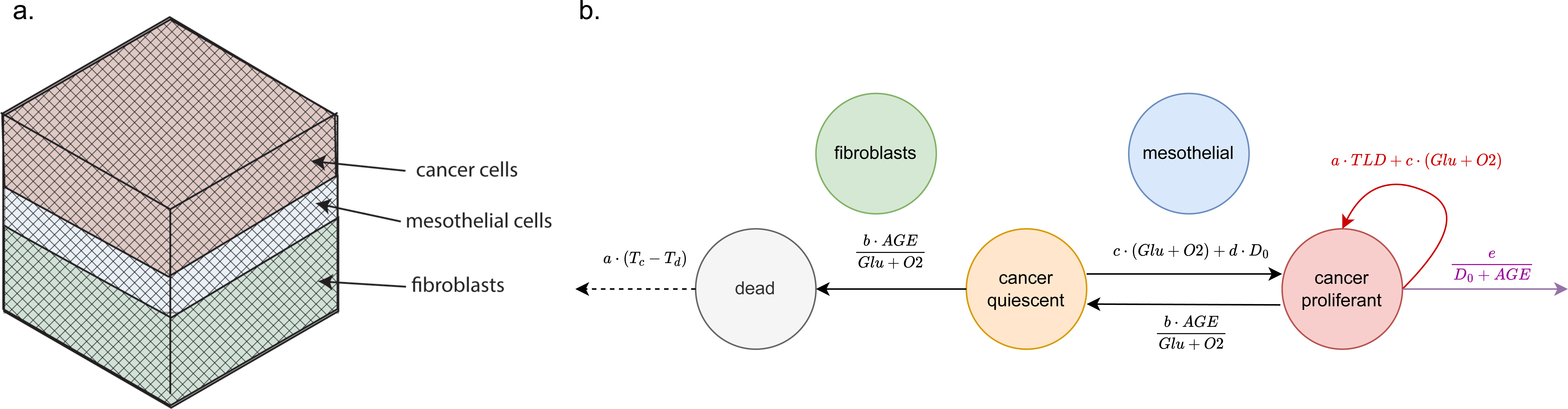
Schematic representation of the SALSA model used within this work. a. Cubic lattice representing the underlying structure of the simulator. Shaded areas distinguish the three main layers of the omentum lining (fibroblast + ECM in green, mesothelial in blue and cancer in red). b. Flowchart of the states (nodes) and behaviours (arcs) formalised within the CM. Beside transitions between different states (black solid arrows) proliferating cancer cells can duplicate (red arrow) and migrate (purple arrow), while dead cells can degrade (dotted arrow). The equations on each arc represent the probability of occurrence of each rule and the definition of the variables they contain is in Tab 1.

Modifications to the SALSA seeding procedure were implemented to replicate the layered structure of the omentum (Fig 1a). Fibroblasts cells were limited to the bottom half of the model, with mesothelial cells located immediately on top. Cancer cells were initially positioned above the mesothelial layer and were constrained to move toward the bottom, as the region above them represents the peritoneal cavity. The position of each cell, within the specific region, was randomly determined as in previous versions of SALSA.

The computational representation of the 3D organotypic model comprises 5 different cell states: (i) fibroblasts, (ii) mesothelial cells (iii) dead PEO4 cells, (iv) quiescent PEO4 cells and (v) proliferating PEO4 cells (Fig 1b). Arcs within the graph in Fig 1b represent the allowed transitions between different states and are labelled with the corresponding probability of occurrence (see Tab 1 for the definition of each variable). These rules were determined collating multiple evidence from the scientific literature, with the aim of describing the behaviour of PEO4 cells. Parameters a-e were empirically estimated through the procedure described in the following section. Fibroblasts and mesothelial cells don’t have a formalised behaviour and are assumed to maintain their status throughout the simulation. They however consume resources (glucose and oxygen), thus impacting indirectly the behaviour of the HGSOC cells.

**Table 1.** Variables used to formalise cell behaviour and their definition

| Variable | Definition |
| --- | --- |
| AGE | Age of the cell normalised with respect to the length of the simulation |
| D | Local drug concentration normalised with respect to the amount added in the media |
| D<sub>0</sub> | Distance between the position of the current cell and the bottom of the culture (normalised between 0 and 1) |
| Glu | Local glucose level normalised with respect to its concentration in the media |
| O2 | Local oxygen level normalised with respect to its concentration in the media |
| T<sub>c</sub> | Current time point normalised with respect to the length of the simulation |
| T<sub>d</sub> | Time at which the current cell died normalised with respect to the length of the simulation |
| TLD | Time elapsed since the last division of the current cell normalised with respect to the length of the simulation |

Treatment with cisplatin and paclitaxel was modelled as described in [45]. A sigmoid curve (S) was used to describe the probability of the drug affecting cell behaviour as a function of the local drug concentration (D). The parameters of this response curve were identified, using the IC_50_ values and assuming no effect in absence of the drug. In both cases, only proliferation (PR) and the rate of cell death (CDR) were considered to be affected by the treatment (see Eqs 1 2,S2 Fig and Tab 1 for definitions), in accordance with the mechanism of action of these agents [47, 48].

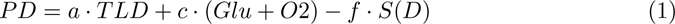

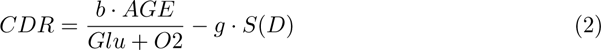

### Parameters Estimation

Initially, the parameters describing the behaviour of the culture in absence of treatment (Fig. 1) were calibrated. A number of different values between 0 and 1 were considered for each parameter (0.001, 0.005, 0.01, 0.05, 0.1, 0.5, 1) and every possible combination of these values was simulated 3 times for the equivalent of 72 h. The replicates were then averaged and a score comparing simulated and *in-vitro* data was computed (Eq. 3).

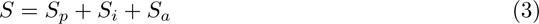

Here S_p_, S_i_ and S_a_ are defined as in Eqs 4 5 6, where C_x_ is the number of simulated cancer cells at T = x. DR is the experimentally measured doubling rate. It was set to 2 for the monolayer cultures (based on a doubling rate of 36 h [36] and to 0.87 for the 3D hydrogel multispheroids. This value was obtained as the number of cells counted in bright-field images obtained during a 7 day long experiment in the IncuCyte real-time cell imaging instrument (S3 Fig a.). Finally, I_silico_ and I_vitro_ are the number of invaded HGSOC cells and A_x_ the measured amount of adherent PEO4 cells at T = x.

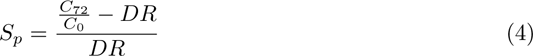

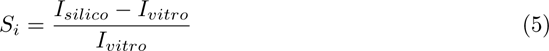

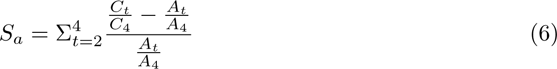

S_p_ measures the effectiveness of the computational model in recapitulating cancer cell proliferation.

S_i_ achieves the same purpose, but it compares the number of invaded cells measured *in-vitro* with the average number of migration events recorded during the simulation.

S_a_ has a similar structure, but it compares the number of simulated PEO4 cells at iterations 2-4 to the corresponding results of the adhesion experiments.

A total of 8 different computational models were identified, using every possible combination of 2D and 3D data (Tab 2) and choosing the parameter set associated with the best (i.e., lowest) score value.

**Table 2.**
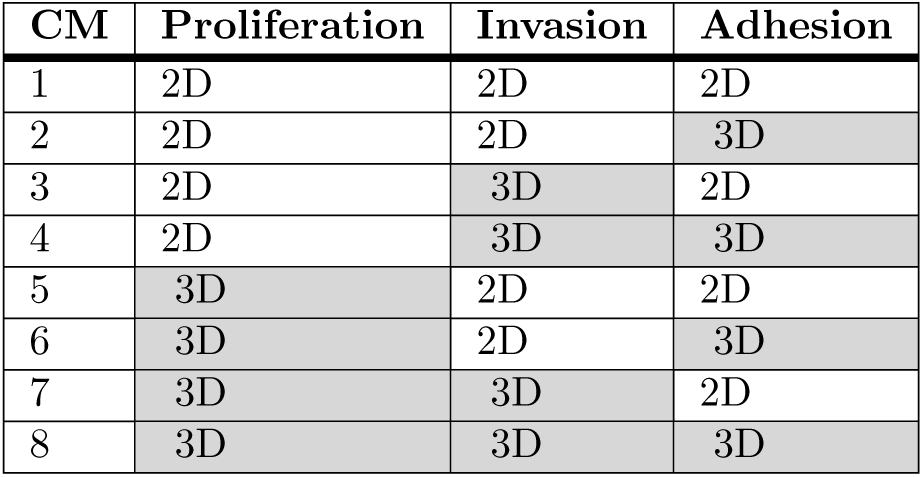
Combination of the 2D (white background) and 3D (gray background) data used for the analysis.

The same procedure was repeated for the two parameters recapitulating the drug treatment. In this case, only the IC_50_ values for cisplatin and paclitaxel (10.4 *µ*M and 3.04 nM respectively, [49]) were used for the calibration, and the configuration corresponding to a normalised cancer cell density closest to 0.5 was selected.

### Computational model validation

The validation of the CMs was conducted by comparing the simulated dose response curves to cisplatin and paclitaxel, with the corresponding experimental data acquired in monolayer cultures and 3D multi-spheroids. No modification to the structure or parameters of the model was applied, with respect to the calibration stage, and the *in-vitro* data used for the comparison were not used to identify any of the parameters.

### Statistical analysis

The Kolmogorov-Smirnov test was used, whenever appropriate, to evaluate whether the distribution underlying the two sets of samples was the same. This method was chosen as it does not make any assumption on the shape of the underlying distribution. A p-value of 0.05 was chosen as the threshold for significance.

## Results

Our analysis is schematically described in Fig 2. We considered three EMs: 3D hydrogel multi-spheroids, a 3D organotypic model and standard monolayer culture. The use of two 3D EMs was required as the CMs describe both the first phases of metastasis, when cancer cells exhibit limited proliferation but readily adhere and invade and the later stages of disease progression when cells proliferate within the invaded tissue. The organotypic model is an accurate representation of the adhesion/invasion phase, but constraints on the length of the experiments and difficulties in separating the contribution of the different cell types limit its usefulness for the evaluation of proliferation. Hydrogel multi-spheroids, on the other hand, yield accurate proliferation measurements, but lack the multilayer structure useful for the study of invasion. As such, we decided to exploit the strengths of both systems and evaluate adhesion and invasion in the organotypic model and proliferation and drug response in the 3D hydrogel multi-spheroids. The same measures were also obtained in standard 2D cell cultures.

**Fig 2.**
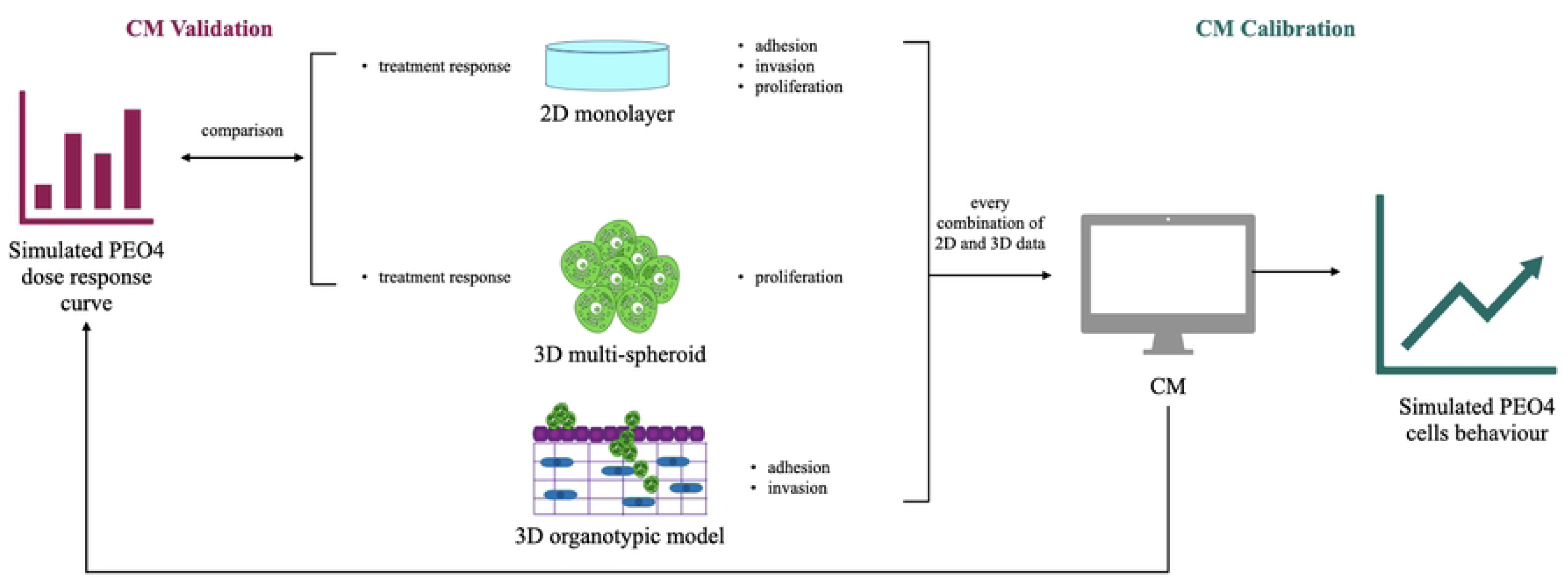
Flowchart of the analysis presented in this work. Different combinations of experimental data from 2D monolayers and 2 3D EMs (hydrogel multi-spheroids and an organotypic model) were used to calibrate the same computational simulator of transcoelomic metastasis. These CMs were then used to simulate the response to either cisplatin or paclitaxel. The comparison between the simulated and measured dose response curves enabled the validation of the CMs and thus the determination of which CM yields the results more closely matching experimental data.

Adhesion, invasion and proliferation data were used to calibrate the CMs (right end side of Fig 2). To test the effect of using different EMs on the simulated results we considered all possible combinations of 2D and 3D data. The calibrated CMs allowed study of the *in-silico* behaviour of PEO4 cells both in absence and presence of treatment. The simulated dose response curves were compared with their experimental counterpart, acquired both in 2D monolayers and in the 3D hydrogel multi-spheroids, to determine which combination of calibration data yielded the CM better replicating the treatment response of PEO4 cells (left end side of Fig 2).

### *in-vitro* quantification of adhesion and invasion

Fig 3a reports the results of the adhesion time course conducted in both 2D monolayers and the organotypic model. Very similar absorbance values were obtained for the two conditions, even though a 3D setting was associated with increased variability. Additionally, the number of adherent cells remained approximately constant within the considered timeframe, even though a slight trend appears to be present for the 3D setting.

**Fig 3.**
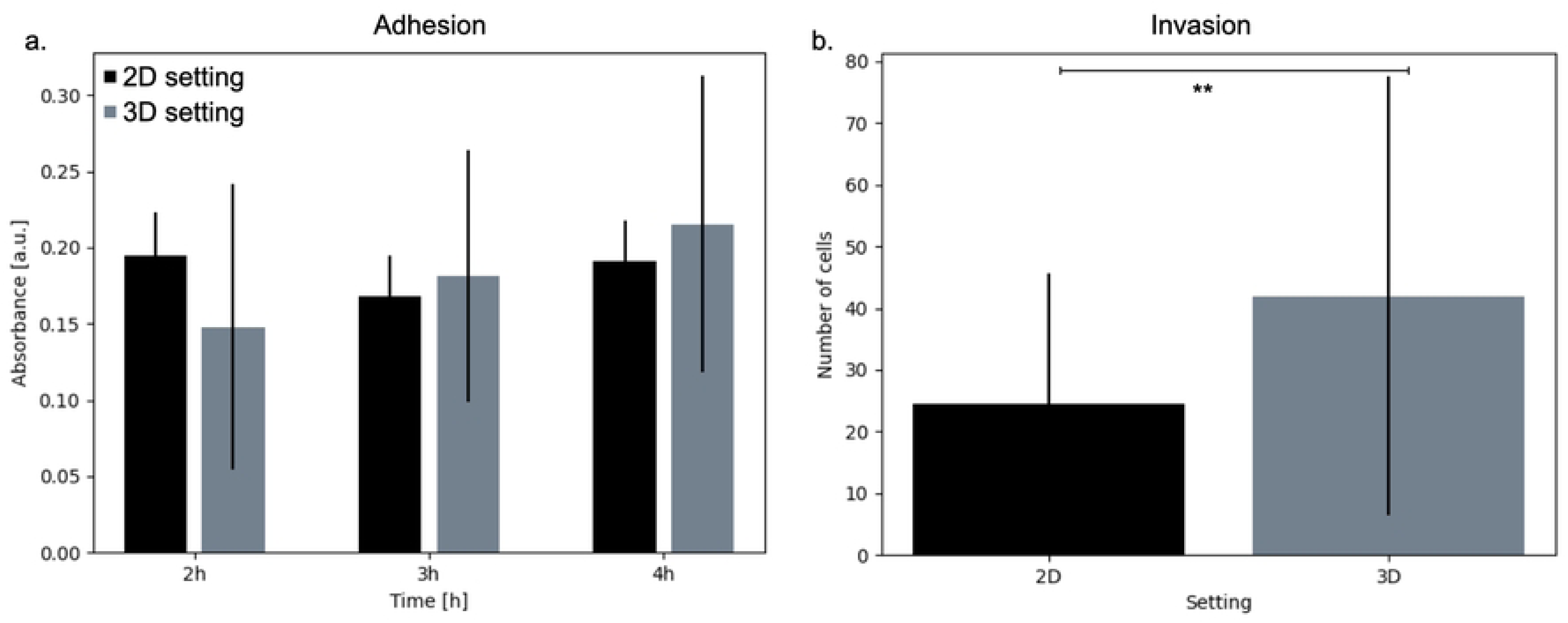
Experimentally measured adhesion and invasion in both 2D and 3D EMs. a. Adhesion measurements at 2, 3 and 4 h post seeding. In 2D a collagen coating was used as substrate while in 3D HGSOC cells were seeded on the organotypic model. b. Average number of invaded PEO4 cells in 2D and 3D (Kolmogorov Smirnov p value = 0.005). In both panels error bars represent the standard deviation (n = 3).

Changing the EM had a more pronounced effect when quantifying invasion (Fig 3b). Here the use of the organotypic model resulted in a noticeable increase in the number of invading cells (Kolmogorov-Smirnov test p = 0.005).

These results, together with the doubling time for PEO4 cells [36] and their confluency measurement obtained with the IncuCyte (S3 Fig), were used to calibrate the CMs, that is to identify which parameter sets better approximate the different combination of experimental data.

### Computational models calibration

To determine the role of the EM in the identification of the CM parameters we calibrated eight different CMs, each corresponding to a different combination of experimental data (Tab 2). As described in the methods section, a score was computed for each simulated parameters configuration (Eq 3), and the one associated with the lowest value was selected.

Fig 4 reports the results of this analysis both as individual score components (panels a. to c.) and overall score value (panel d.). For each CM the score values are reported as average and standard deviation (computed over 50 simulations). No statistical analysis was conducted on these data, as the accuracy of each CM in reproducing the experimental results was assessed independently.

**Fig 4.**
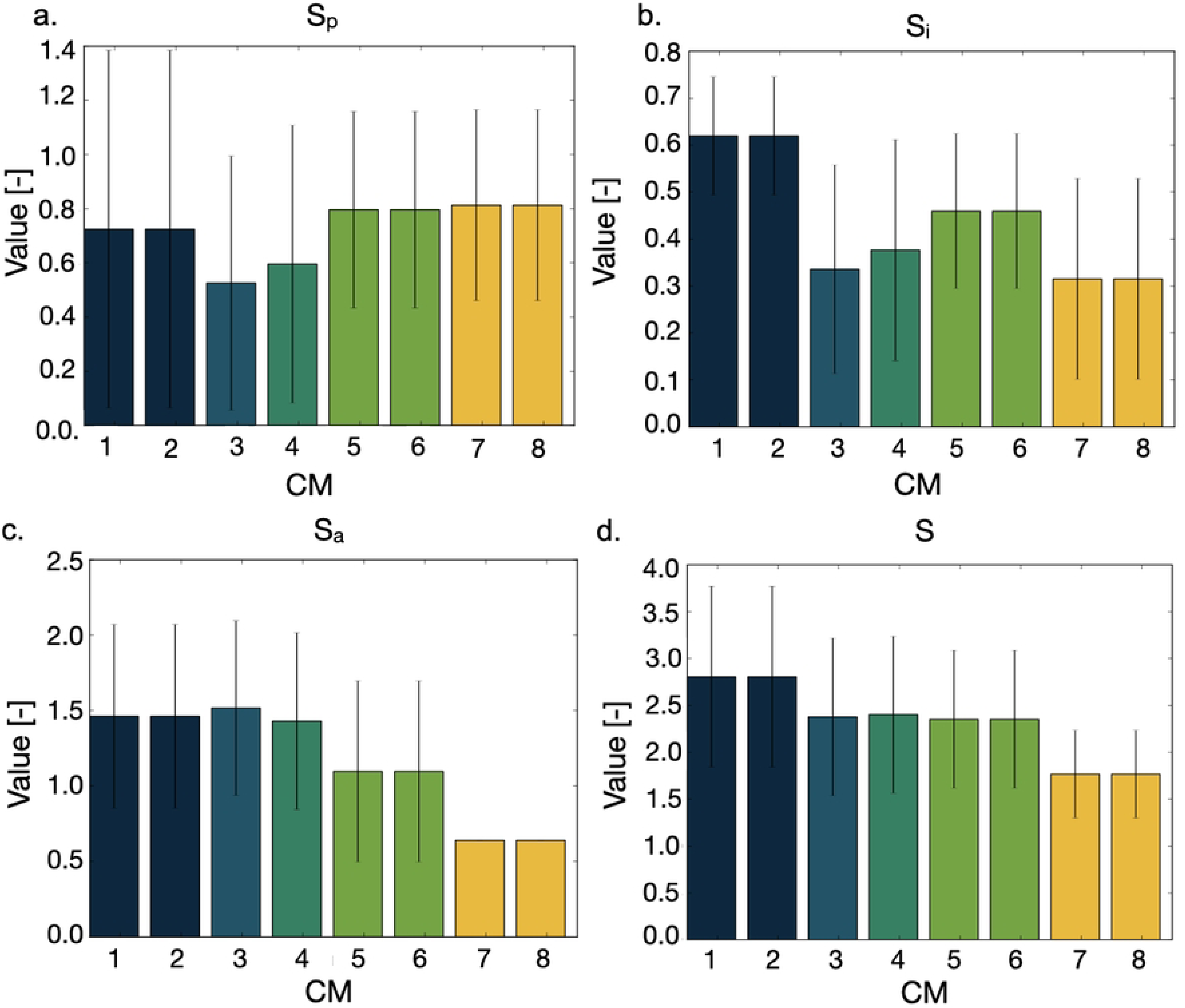
Score values associated with the best parameter configuration for each CM. Panels a-c refer to one of the three score components (a. S_p_, b. S_i_, c. S_a_), while panel d. shows each configuration overall score. As-per their definition (see Eqs 3, 4, 5, 6) low score values are associated with higher accuracy of the CM. Scores were computed independently for each simulation and then averaged (error bars represent the standard deviation). The experimental data used to calibrate each CM are summarised in Tab. 2 and the corresponding colour coding is reported in Tab 3

The use of experimental data acquired in 3D seems to be associated with an overall lower score (1.7 for CM 8 vs 2.8 for CM 1), but the main result of this analysis is the low relevance of the EM used to evaluate adhesion. Indeed, in most cases, both 2D and 3D experimental data yield the same parameter set (Tab 3). The only exceptions are CMs 3 and 4, which are however associated with very similar configurations (Tab 3) and overall score (Fig 4).

**Table 3.** Optimal parameter configurations for each CM in Tab. 2 CMs with identical parameters values (e.g. 1 and 2) will be considered as one condition (1/2) for the rest of the analysis. Each CM has also been color-coded (last column of the table) throughout the analysis

| CM |  | Parameters |  |  |  |  | Colour |
| --- | --- | --- | --- | --- | --- | --- | --- |
|  |  | a | b | c | d | e |  |
| 1<br>2 | 1/2 | 1 | 0.1 | 0.1 | 0.01 | 0.1 |  |
| 3 |  | 1 | 0.5 | 0.1 | 0.01 | 0.1 |  |
| 4 |  | 1 | 0.5 | 0.1 | 0.1 | 0.1 |  |
| 5<br>6 | 5/6 | 1 | 0.1 | 0.05 | 0.001 | 0.5 |  |
| 7<br>8 | 7/8 | 1 | 0.5 | 0.05 | 1 | 0.5 |  |

The parameters describing drug response were determined in the same way. Tab 4 reports the values of f and g (Eqs 1 2) for each CM and the corresponding score value. Again, the error associated with the use of 3D data is generally lower, even though no general association between use of specific EMs and parameter values was observed.

**Table 4.** Optimal parameter configurations for treatment response simulation for each CM.

| CM | Cisplatin |  |  | Paclitaxel |  |  |
| --- | --- | --- | --- | --- | --- | --- |
|  | f | g | score | f | g | score |
| 1/2 | 0.5 | 0.01 | 1.2 | 0.05 | 0.001 | 1.42 |
| 3 | 0.5 | 0.5 | 0.16 | 0.005 | 0.5 | 0.05 |
| 4 | 1 | 0.005 | 0.26 | 1 | 0.01 | 0.37 |
| 5/6 | 0.05 | 0.001 | 0.16 | 0.005 | 0.01 | 0.16 |
| 7/8 | 1 | 0.01 | 0.05 | 0.01 | 0.005 | 0.05 |

### Computational model validation

Following the identification of the CMs, we compared the simulated response to treatment with either cisplatin (Fig 5) or paclitaxel (Fig. 6) to the experimentally measured values in a 2D or 3D setting. Most of the CMs showed limited response to treatment and a statistically significant difference between the simulated and experimental data (Kolmogorov-Smirnov test, * p *<*0.05, ** p*<*0.01), especially at the higher drug concentrations.

**Fig 5.**
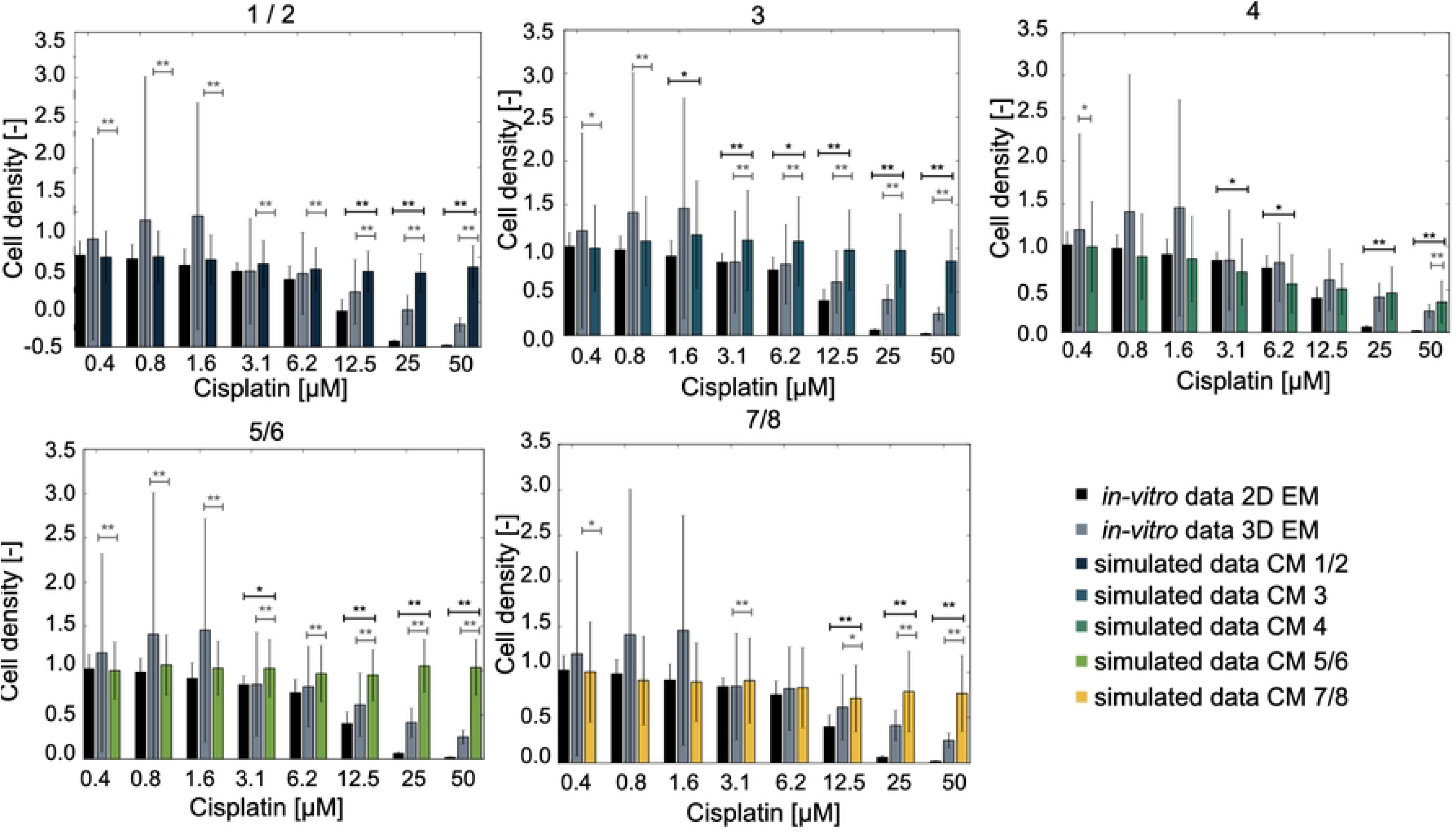
Comparison between the simulated cisplatin response (colour coded for each model as in Table 3) and the experimental data acquired in 2D (red bars) and 3D (purple bars) EMs. In all cases, the cell density is normalised with respect to untreated condition and data are reported as mean +/- standard deviation (n=3 for the experimental data, n=50 for the simulated results) Statistical testing conducted using the Kolmogorov-Smirnov test and reported in black when comparing simulated and 2D data and in grey for simulated and 3D data, * p*<*0.05, **p*<*0.01.

**Fig 6.**
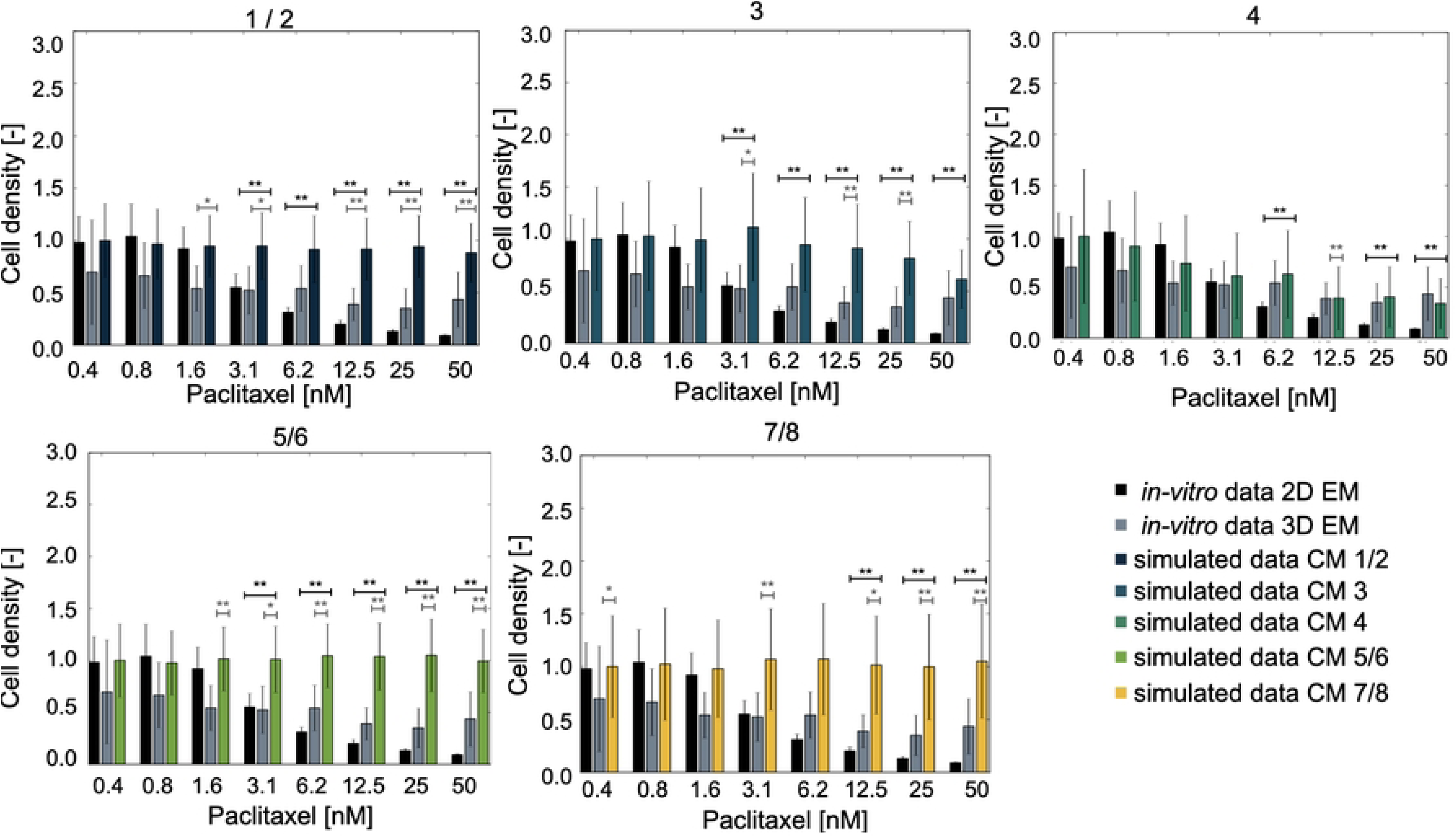
Comparison between the simulated paclitaxel response (colour coded for each model as in Table 3) and the experimental data acquired in 2D (red bars) and 3D (purple bars) EMs. In all cases, the cell density is normalised with respect to untreated condition and data are reported as mean +/- standard deviation (n=3 for the experimental data, n=50 for the simulated results) Statistical testing conducted using the Kolmogorov-Smirnov test and reported in black when comparing simulated and 2D data and in grey for simulated and 3D data, * p*<*0.05, **p*<*0.01.

Only CM 4 recapitulated the dose response curve accurately. It corresponds to using 3D data for both invasion and adhesion measurements and 2D measurements for proliferation (Tab. 2). This result is also confirmed by relative error between experimental and simulated data (Tab. 5). In this case the relative change between experimental and simulated results was computed for each drug concentration, and then summed to provide an indication of how well the model captures the entire dose response curve.

**Table 5.** Optimal parameter configurations for treatment response simulation for each CM.

| CM | Cisplatin |  | Paclitaxel |  |
| --- | --- | --- | --- | --- |
|  | 2D | 3D | 2D | 3D |
| 1/2 | 59.29 | 4.88 | 21.42 | 7.21 |
| 3 | 60.89 | 5.61 | 17.84 | 6.78 |
| 4 | **24.81** | **2.13** | **7.33** | **1.85** |
| 5/6 | 71.34 | 6.36 | 24.70 | 8.59 |
| 7/8 | 51.82 | 4.14 | 18.71 | 4.92 |

3D data tend to be better approximated by the computational model, as the error is lower. This might be partly due to the more limited growth rate and response to treatment observed in 3D, which might be simpler to capture with the simulator. At the same time, this difference between treatment in 2D and 3D settings is commonly observed, both in terms of increased IC_50_ and reduced overall response [50, 51].

### High resolution analysis of simulated cell behaviour

A key feature of SALSA is that it retains information on the position of each cell at each iteration. This enables study of the dynamic distribution of each population with sub-organoid resolution. In particular, the distribution of the average cell density for each simulated cell type was computed as a function of time (x axis) and z coordinate (y axis). Cell density was obtained, for each simulation, normalising the number of cells at each depth by the initial population cardinality. Averaging over the simulation yielded the heatmaps in Figs. 7, 8, 9.

**Fig 7.**
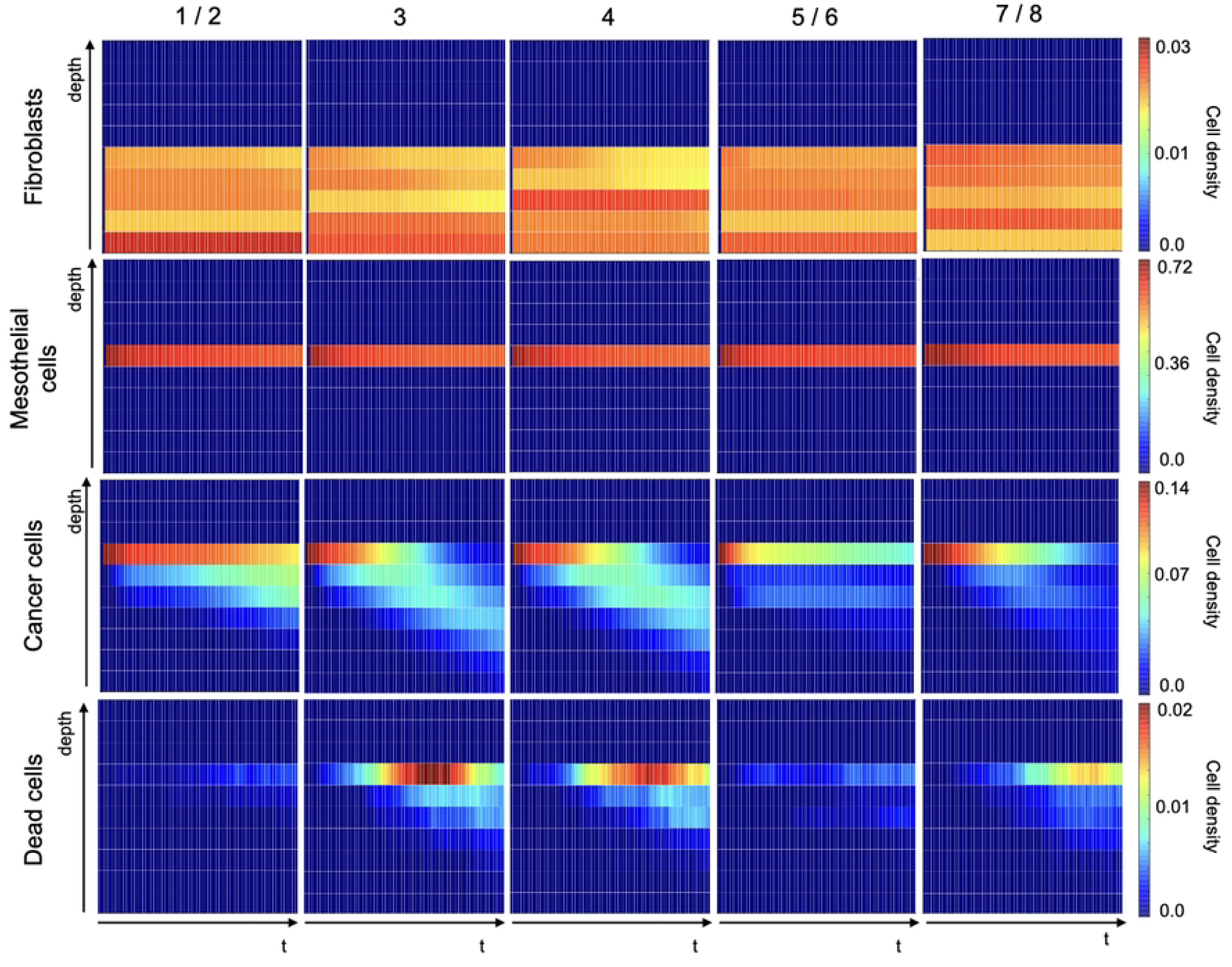
Analysis of the dynamic behaviour of simulated cell types with sub-organoid resolution. Each column corresponds to a different CM, while each row is associated to a cell type. Every panel shows the average density of that cell type (over 50 simulations) over time and as a function of the z coordinate. Colour shading in each block represents cell density (refer to scale on right end side), with dark blue and bright red representing the lower and higher cell densities. All values have been normalised with respect to total initial cell number.

**Fig 8.**
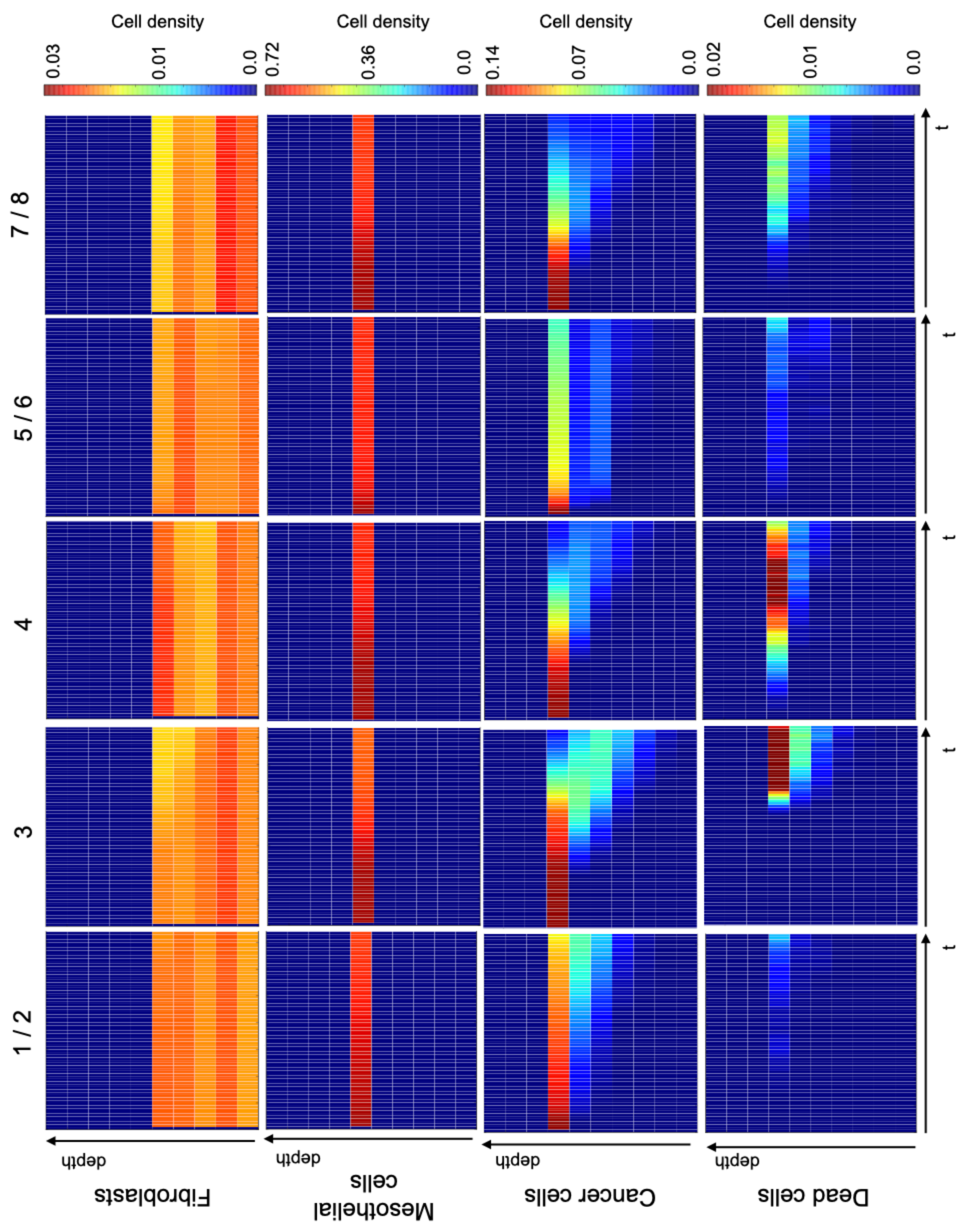
Analysis of the dynamic behaviour of simulated cell types with sub-organoid resolution following treatment with cisplatin at IC_50_ levels. Each column corresponds to a different CM, while each row is associated to a cell type. Every panel shows the average density of that cell type (over 50 simulations) over time and as a function of the z coordinate. Colour shading in each block represents cell density (refer to scale on right end side), with dark blue and bright red representing the lower and higher cell densities. All values have been normalised with respect to total initial cell number.

**Fig 9.**
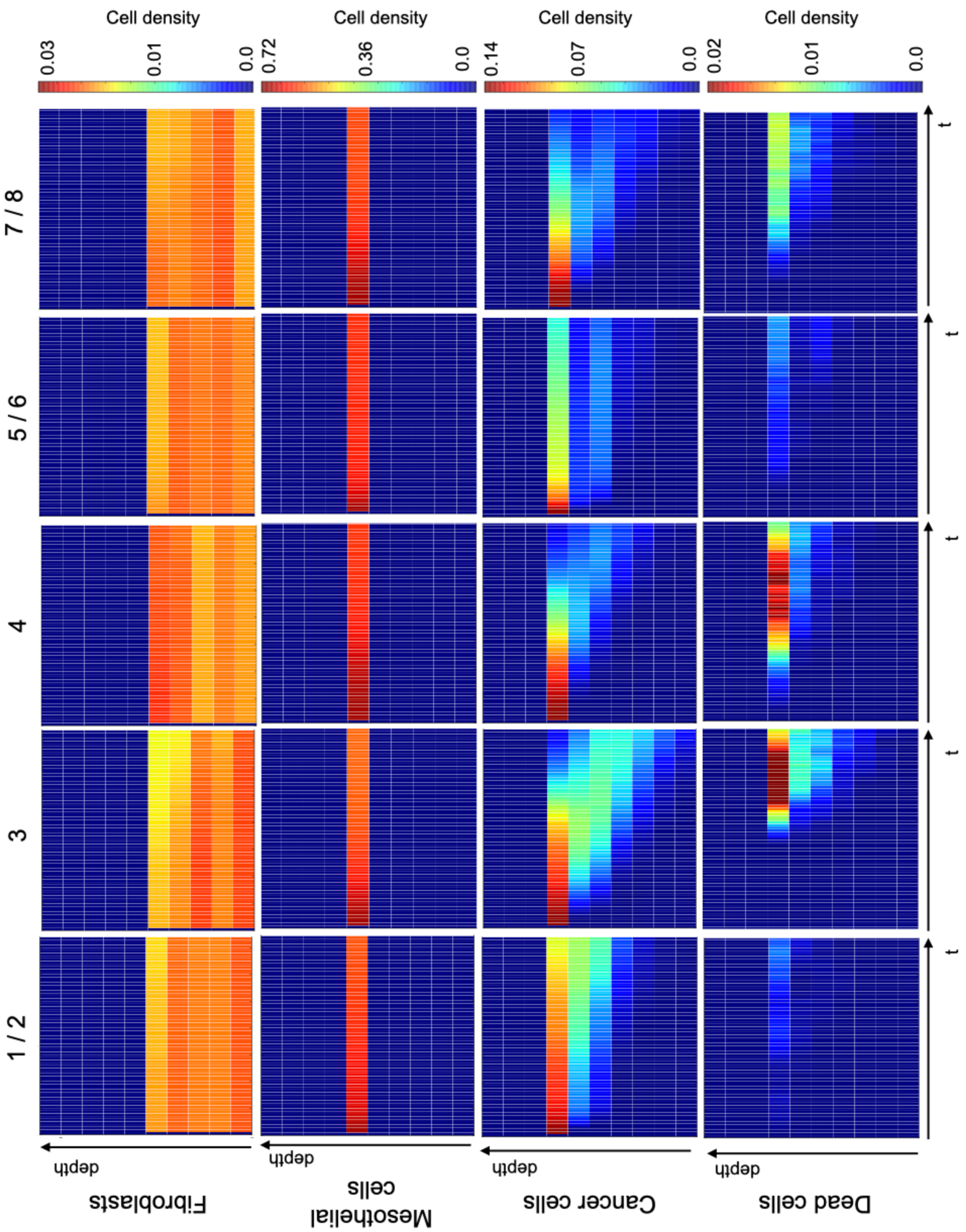
Analysis of the dynamic behaviour of simulated cell types with sub-organoid resolution following treatment with paclitaxel at IC_50_ levels. Each column corresponds to a different CM, while each row is associated to a cell type. Every panel shows the average density of that cell type (over 50 simulations) over time and as a function of the z coordinate. Colour shading in each block represents cell density (refer to scale on right end side), with dark blue and bright red representing the lower and higher cell densities. All values have been normalised with respect to total initial cell number.

The difference between them is the treatment condition: Fig. 7 outlines the behaviour of the different CMs in the absence of treatment while Figs. 8 and 9 refer to treatment with cisplatin and paclitaxel at a concentration equal to the simulated IC_50_. Fibroblast and mesothelial cells are concentrated in very specific regions of the virtual organoid, in accordance with the definition of the experimental model [28, 29] and their density is constant throughout the experiment, as per CM definition (Fig. 1). Any variation in the colour shade is due to the fact that virtual cells are randomly assigned a position within the allowed area, and this will result in a non-uniform distribution even when the average density is computed. Additionally, as virtual PEO4 cells move through the organoid they have the power of displacing fibroblasts and mesothelial cells, hence producing the slight decrease in the density of these cells observed as time progresses.

The behaviour of virtual PEO4 cells is conceptually the same in all models: their initial location is on top of the mesothelial cells, and they progressively infiltrate the underlying layers. The different CMs are however characterised by varying degrees of infiltration and proliferation within the organoid.

Configurations calibrated with the 3D invasion data (CMs 3, 4, 7/8) result in a more extensive infiltration (i.e., cancer cells present at lower depth values) while 2D proliferation (CMs 1/2, 3 and 4) is associated with an overall higher number of cancer cells. 3D invasion is also connected with a higher rate of PEO4 cell death especially in the shallower layers of the organoid and the second half of the simulation. This might be connected to the dependence of cell death on the age of the cancer cells, which is likely to be higher toward the end of the simulation and in the region where this type of cells was initially located.

Comparing Fig. 7 with Figs. 8 and 9 highlights the effect of the treatments on the behaviours of the different cell types.

Simulated fibroblasts and mesothelial cells are not affected by the treatment, as their interaction with cisplatin and paclitaxel has not been formalised. PEO4 cells, on the other hand, have a generally reduced growth and more limited infiltration within the virtual organotypic model. This seems to be connected to a delay in both these processes, whose probability increases as the effective drug concentration decreases due to degradation.

Cell death is also increased in the presence of treatment and this process seems to be more sustained over time. Surprisingly however, the number of dead cells becomes relevant at about the same time as in the untreated condition, suggesting that, in our simulations, treatment alone is not sufficient to induce cell quiescence and death. This is consistent with the treatment resistant nature of PEO4 cells but might also reflect the need for a more detailed modelling of drug response.

## 1 Discussion

Computational modelling has acquired great relevance in biomedical and cancer research, both as a tool for fundamental research [52–54] and as an aid for clinical decision making [55]. As such, an in-depth analysis of the calibration and validation procedures is necessary to maximise the utility and accuracy of these models.

The focus of this work has been the effect of using different EMs to calibrate and/or validate the computational simulator and how this choice affects the *in-silico* results. To this end, we measured ovarian cancer cell proliferation, adhesion and invasion in both 2D monolayers and 3D cultures and evaluated the consequences of using distinct combinations of these data for the corroboration of the same computational system, a virtual representation of transcoelomic metastasis realised in SALSA (Fig 1). This biological process was chosen as it is highly dependent on the 3D interaction between different kinds of the cells and their environment, a feature expected to magnify the difference between more accurate experimental models and simplified systems.

The use of different datasets led to the identification of CMs characterised by distinct parameters (Tabs 3, 4) and a varying degree of accuracy when compared with the experimental data (Fig 4, Tab. 4). From these results, the use of at least some of data acquired in a 3D setting tends to be associated with a lower error, underscoring the importance of accurate *in-vitro* models for the corroboration of *in-silico* systems. It is also worth noticing the dependence of some rates on specific experimental data.

The value of the b parameter, which modulates the death rate and the transition between proliferative and quiescent states (Tab. 3), is strictly associated with the experimental model used to evaluate invasion, suggesting that culture in the organotypic model increases the likelihood of both these phenomena. On the other hand, the values of c and e (Tab. 3) are mainly determined by the setting used to measure proliferation. As such, 2D monolayer seems to favour proliferation, while growing in 3D promotes migration and invasion. This is qualitatively coherent with with evidence from the literature [35] and is also confirmed by our experimental data, as the average number of cells able to invade through the organotypic model is almost double that measured in the 2D setting (Fig. 3b). These considerations are also supported by the results presented in Fig 7, where simulated dynamic evolution of the density of the different cell types is presented as a function of their position within the organotypic model. CMs calibrated with 3D invasion data (CMs 3, 4, 7 / 8) are associated with a deeper infiltration of cancer cells and a higher density of dead cells. 3D proliferation (CMs 5/6 and 7/8), on the other hand results in an overall lower number of cancer cells, in agreement with the *in-vitro* data used for the calibration.

The EM used to evaluate cell adhesion seems to be less relevant with 6 out of 8 models being invariant with respect to this property. This could be due to the similarity between the measurements obtained in the two experimental setups (Fig. 3a) but is also a reflection of the mainly surface nature of this phenomenon, which might resent less from the simplification in the experimental setting.

Another key difference between the experimental data measured in 2D and 3D is the increase in variability in the latter (Figs. 3, 5, 6). This is a phenomenon frequently observed when transitioning from monolayer cultures to more complex setups which has been linked to differences in the microenvironments experienced by each cell and have been shown to improve resilience and adaptability [56].

The validation of these CMs was conducted by comparing the simulated dose response curves to cisplatin and paclitaxel with the corresponding experimental results obtained in 2D monolayers and 3D hydrogel models (Figs 5, 6). Data acquired in the multi-spheroid model are associated with a reduced response to the treatments (gray vs black bars in Figs 5, 6) and with an increase in variability. Most CMs are also associated with a limited response to treatment and a low sensitivity to the change in drug concentration. This is likely dependent on the formalization of cancer cell behaviour and response to treatment used within this work, and further analysis on how changing the CM structure and probabilities affects the simulated results is warranted. Beside potentially improving the simulated drug response, this perspective study would also shed light on the role of each simulated variable (e.g., nutrients availability, cell age) in determining cell behaviour, and could provide useful insights on the biology of HGSOC cells. One of the simulated CMs (CM 4), however, was able to effectively recapitulate 2D and 3D dose response curves. It was calibrated using 2D proliferation data and 3D invasion and adhesion measurements. Furthermore, the drug response parameters for this configuration feature a high rate of proliferation inhibition and a comparatively low induction of cell death. Overall, these characteristics produce a response comparable to model 8, which was calibrated using only 3D data, at low and mid drug levels (Figs 5, 6, 8, 9). It however produces a better response when higher concentrations of treatment are simulated. Particularly relevant is the comparison with CM 3, which exhibited a substantially equivalent behaviour in absence of treatment (Fig. 4) but a much more drug-resistant phenotype (Figs 5, 6). The values of parameters f and g for these configurations suggest that, at least in this configuration, inhibiting cell proliferation might be a more effective treatment strategy than inducing cell death.

## Conclusion

Overall, while the results of this work might not be directly transferrable to different experimental and computational models, this analysis allows to draw three conclusions with general applicability.

Firstly, the interaction of multiple phenomena can result in different datasets producing similar results. This can be observed in Figs 8, 9, where CMs 4 and 7 / 8 yield comparable cancer cells densities, despite having been calibrated with different datasets. In this case, a higher number of dead cells in CM 4 compensates for the faster proliferation of this configuration. The comparative analysis presented in this work can be a useful tool for the identification of these compensatory mechanisms, thus enabling a deeper understanding of the computational models and their mechanisms.

Secondly, maximising the similarity between the simulated and experimental setups is associated with a low error throughout all the analysis. Indeed, while a combination of 2D and 3D data most accurately captured the response to cisplatin and paclitaxel, exclusively using data measured in 3D settings scored the lowest in the calibration stage (Fig 3) and second lowest in the drug response (Tab 4, 5).

Thirdly, behaviours more strictly connected with the interaction between the cells and the 3D environment seem to be affected more deeply by simplifications of the experimental model. In our experiments, the number of invaded cells almost doubled shifting from a 2D to a 3D setting (Fig 3b.) and the behaviour of models corroborated with 3D invasion data was markedly different from their 2D counterpart. On the other hand, quantifying adhesion in a 2D or 3D setting had little effect on the CM’s results. As such, should it not be possible to use exclusively 3D EMs, properties characterised by limited cell-cell and cell-environment interactions are expected to be the less affected by the simplification of the EM.

Overall, many obstacles are still in the way of a complete integration of *in-silico* and *in-vitro* analyses, but this and other works provide important insights on how these issues can be addressed and workflows adapted to reap the benefits of computational analysis for the study of complex biological processes.

## Supporting information

**S1 Fig. Quantification of the number of invaded cells** a. Representative image of the DAPI stained nuclei of cancer cells during a 2D invasion assay. b. Segmentation of the image in a., individual cells are represented with different colours to distinguish them. c. Comparison between automatic and manual counts for all the images acquired during the 2D experiments (R2 =0.95). d. Representative image of the DAPI stained nuclei during a 3D invasion assay. e. Segmentation of the image in d.. In this case, additional filters on the area and the GFP intensity are applied to isolate only the cancer cells (colour overlays). f. GFP intensity for the image in d..

**S2 Fig. Flowchart showing the modifications introduced in the computational model to describe treatment with cisplatin and paclitaxel.** As mentioned in the text, only the probability of cell doubling and that of cell death are affected.

**S3 Fig. PEO4 cells growth within 3D printed hydrogels.** a. Dynamic evolution of the number of cells over the course of 7 days. For this experiment 3 different initial cell densities were considered (2000, 3000 and 4000 cells/spheroid) and their growth was monitored with a time resolution of 1 h with an IncuCyte device. The time course starts at T = 72 h to avoid any artefact from bubbles released from the hydrogel. b. Cell density in the same samples was also evaluated with a Cell-titer Glo 3D assay at the end of the experiment. As further detailed in the method section, the two methods produce equivalent results.

**S1 File. Configuration file used to describe the organotypic model in SALSA.**

## Supporting information

Supplementary data (SALSA configuration file)

## Acknowledgments

The authors are grateful to A/Prof. Emanuele Giordano for the helpful feedbacks and discussions. They also acknowledge the individuals that donated tissues for our research.

## References

1. Sadria M, Layton AT. Interactions among mTORC, AMPK and SIRT: a computational model for cell energy balance and metabolism. Cell Communication and Signaling. 2021;19(1):1–17.

2. Cortesi M, Pasini A, Furini S, Giordano E. Identification via numerical computation of transcriptional determinants of a cell phenotype decision making. Frontiers in genetics. 2019;10:575.

3. Yuan B, Shen C, Luna A, Korkut A, Marks DS, Ingraham J, et al. CellBox: interpretable machine learning for perturbation biology with application to the design of cancer combination therapy. Cell systems. 2021;12(2):128–140.

4. Zhao C, Medeiros TX, Sové RJ, Annex BH, Popel AS. A data-driven computational model enables integrative and mechanistic characterization of dynamic macrophage polarization. Iscience. 2021;24(2):102112.

5. Shine JM, Müller EJ, Munn B, Cabral J, Moran RJ, Breakspear M. Computational models link cellular mechanisms of neuromodulation to large-scale neural dynamics. Nature neuroscience. 2021;24(6):765–776.

6. Telmer CA, Sayed K, Butchy AA, Bocan K, Kaltenmeier C, Lotze M, et al. Computational modeling of cell signaling and mutations in pancreatic cancer. bioRxiv. 2021; p. 2021–06.

7. Linden NJ, Kramer B, Rangamani P. Bayesian parameter estimation for dynamical models in systems biology. PLOS Computational Biology. 2022;18(10):e1010651.

8. Reali F, Priami C, Marchetti L. Optimization algorithms for computational systems biology. Frontiers in Applied Mathematics and Statistics. 2017;3:6.

9. Schmiester L, Schälte Y, Bergmann FT, Camba T, Dudkin E, Egert J, et al. PEtab—Interoperable specification of parameter estimation problems in systems biology. PLoS computational biology. 2021;17(1):e1008646.

10. Viceconti M, Pappalardo F, Rodriguez B, Horner M, Bischoff J, Tshinanu FM. In silico trials: Verification, validation and uncertainty quantification of predictive models used in the regulatory evaluation of biomedical products. Methods. 2021;185:120–127.

11. Wodarz D, Komarova N. Towards predictive computational models of oncolytic virus therapy: basis for experimental validation and model selection. PloS one. 2009;4(1):e4271.

12. Al Ameri W, Ahmed I, Al-Dasim FM, Ali Mohamoud Y, Al-Azwani IK, Malek JA, et al. Cell type-specific TGF-*β* mediated EMT in 3D and 2D models and its reversal by TGF-*β* receptor kinase inhibitor in ovarian cancer cell lines. International journal of molecular sciences. 2019;20(14):3568.

13. Liu M, Zhang X, Long C, Xu H, Cheng X, Chang J, et al. Collagen-based three-dimensional culture microenvironment promotes epithelial to mesenchymal transition and drug resistance of human ovarian cancer in vitro. RSC advances. 2018;8(16):8910–8919.

14. Tofani LB, Abriata JP, Luiz MT, Marchetti JM, Swiech K. Establishment and characterization of an in vitro 3D ovarian cancer model for drug screening assays. Biotechnology Progress. 2020;36(6):e3034.

15. Yousefi M, Dehghani S, Nosrati R, Ghanei M, Salmaninejad A, Rajaie S, et al. Current insights into the metastasis of epithelial ovarian cancer-hopes and hurdles. Cellular Oncology. 2020;43:515–538.

16. Al Habyan S, Kalos C, Szymborski J, McCaffrey L. Multicellular detachment generates metastatic spheroids during intra-abdominal dissemination in epithelial ovarian cancer. Oncogene. 2018;37(37):5127–5135.

17. Capellero S, Erriquez J, Battistini C, Porporato R, Scotto G, Borella F, et al. Ovarian cancer cells in ascites form aggregates that display a hybrid epithelial-mesenchymal phenotype and allows survival and proliferation of metastasizing cells. International Journal of Molecular Sciences. 2022;23(2):833.

18. Etzerodt A, Moulin M, Doktor TK, Delfini M, Mossadegh-Keller N, Bajenoff M, et al. Tissue-resident macrophages in omentum promote metastatic spread of ovarian cancer. Journal of Experimental Medicine. 2020;217(4).

19. Ford CE, Werner B, Hacker NF, Warton K. The untapped potential of ascites in ovarian cancer research and treatment. British Journal of Cancer. 2020;123(1):9–16.

20. Moss NM, Barbolina MV, Liu Y, Sun L, Munshi HG, Stack MS. Ovarian cancer cell detachment and multicellular aggregate formation are regulated by membrane type 1 matrix metalloproteinase: a potential role in Ip metastatic dissemination. Cancer research. 2009;69(17):7121–7129.

21. Pease JC, Brewer M, Tirnauer JS. Spontaneous spheroid budding from monolayers: a potential contribution to ovarian cancer dissemination. Biology open. 2012;1(7):622–628.

22. Steinkamp MP, Winner KK, Davies S, Muller C, Zhang Y, Hoffman RM, et al. Ovarian tumor attachment, invasion, and vascularization reflect unique microenvironments in the peritoneum: insights from xenograft and mathematical models. Frontiers in oncology. 2013;3:97.

23. Borghese C, Casagrande N, Corona G, Aldinucci D. Adipose-derived stem cells primed with paclitaxel inhibit ovarian cancer spheroid growth and overcome paclitaxel resistance. Pharmaceutics. 2020;12(5):401.

24. Braccini S, Tacchini C, Chiellini F, Puppi D. Polymeric Hydrogels for In Vitro 3D Ovarian Cancer Modeling. International Journal of Molecular Sciences. 2022;23(6):3265.

25. Ciucci A, Buttarelli M, Fagotti A, Scambia G, Gallo D. Preclinical models of epithelial ovarian cancer: practical considerations and challenges for a meaningful application. Cellular and Molecular Life Sciences. 2022;79(7):364.

26. Hedegaard CL, Redondo-Gómez C, Tan BY, Ng KW, Loessner D, Mata A. Peptide-protein coassembling matrices as a biomimetic 3D model of ovarian cancer. Science Advances. 2020;6(40):eabb3298.

27. Yee C, Dickson KA, Muntasir MN, Ma Y, Marsh DJ. Three-dimensional modelling of ovarian cancer: From cell lines to organoids for discovery and personalized medicine. Frontiers in Bioengineering and Biotechnology. 2022; p. 116.

28. Kenny HA, Lal-Nag M, White EA, Shen M, Chiang CY, Mitra AK, et al. Quantitative high throughput screening using a primary human three-dimensional organotypic culture predicts in vivo efficacy. Nature communications. 2015;6(1):6220.

29. Peters PN, Schryver EM, Lengyel E, Kenny H. Modeling the early steps of ovarian cancer dissemination in an organotypic culture of the human peritoneal cavity. Journal of visualized experiments: JoVE. 2015;(106).

30. Hart PC, Bajwa P, Kenny HA. Modeling the Early Steps of Ovarian Cancer Dissemination in an Organotypic Culture of the Human Peritoneal Cavity. Ovarian Cancer: Molecular & Diagnostic Imaging and Treatment Strategies. 2021; p. 75–94.

31. Henry C, Hacker N, Ford C. Silencing ROR1 and ROR2 inhibits invasion and adhesion in an organotypic model of ovarian cancer metastasis. Oncotarget. 2017;8(68):112727.

32. Joshi N, Liu D, Dickson KA, Marsh DJ, Ford CE, Stenzel MH. An organotypic model of high-grade serous ovarian cancer to test the anti-metastatic potential of ROR2 targeted Polyion complex nanoparticles. Journal of Materials Chemistry B. 2021;9(44):9123–9135.

33. Kenny HA, Lal-Nag M, Shen M, Kara B, Nahotko DA, Wroblewski K, et al. Quantitative high-throughput screening using an organotypic model identifies compounds that inhibit ovarian cancer metastasis. Molecular cancer therapeutics. 2020;19(1):52–62.

34. Watters KM, Bajwa P, Kenny HA. Organotypic 3D models of the ovarian cancer tumor microenvironment. Cancers. 2018;10(8):265.

35. Kumari A, Shonibare Z, Monavarian M, Arend RC, Lee NY, Inman GJ, et al. TGF*β* signaling networks in ovarian cancer progression and plasticity. Clinical & Experimental Metastasis. 2021;38:139–161.

36. Langdon SP, Lawrie SS, Hay FG, Hawkes MM, McDonald A, Hayward IP, et al. Characterization and properties of nine human ovarian adenocarcinoma cell lines. Cancer research. 1988;48(21):6166–6172.

37. Ng CK, Cooke SL, Howe K, Newman S, Xian J, Temple J, et al. The role of tandem duplicator phenotype in tumour evolution in high-grade serous ovarian cancer. The Journal of pathology. 2012;226(5):703–712.

38. Lu M, Henry CE, Lai H, Khine YY, Ford CE, Stenzel MH. A new 3D organotypic model of ovarian cancer to help evaluate the antimetastatic activity of RAPTA-C conjugated micelles. Biomaterials science. 2019;7(4):1652–1660.

39. Jung M, Skhinas JN, Du EY, Tolentino MK, Utama RH, Engel M, et al. A high-throughput 3D bioprinted cancer cell migration and invasion model with versatile and broad biological applicability. Biomaterials Science. 2022;10(20):5876–5887.

40. Utama RH, Tan VT, Tjandra KC, Sexton A, Nguyen DH, O’Mahony AP, et al. A Covalently Crosslinked Ink for Multimaterials Drop-on-Demand 3D Bioprinting of 3D Cell Cultures. Macromolecular Bioscience. 2021;21(9):2100125.

41. Mauri E, Sacchetti A, Rossi F. The synthesis of RGD-functionalized hydrogels as a tool for therapeutic applications. JoVE (Journal of Visualized Experiments). 2016;(116):e54445.

42. Alday-Parejo B, Ghimire K, Coquoz O, Albisetti GW, Tamò L, Zaric J, et al. MAGI1 localizes to mature focal adhesion and modulates endothelial cell adhesion, migration and angiogenesis. Cell Adhesion & Migration. 2021;15(1):126–139.

43. Cortesi M, Llamosas E, Henry CE, Kumaran RYA, Ng B, Youkhana J, et al. I-AbACUS: a reliable software tool for the semi-automatic analysis of invasion and migration transwell assays. Scientific Reports. 2018;8(1):3814.

44. Cortesi M, Liverani C, Mercatali L, Ibrahim T, Giordano E. An in-silico study of cancer cell survival and spatial distribution within a 3D microenvironment. Scientific Reports. 2020;10(1):1–14.

45. Cortesi M, Liverani C, Mercatali L, Ibrahim T, Giordano E. Development and validation of an in-silico tool for the study of therapeutic agents in 3D cell cultures. Computers in Biology and Medicine. 2021;130:104211.

46. Cortesi M, Giordano E. Driving cell response through deep learning, a study in simulated 3D cell cultures. Heliyon. under review;-(-):–.

47. Dasari S, Tchounwou PB. Cisplatin in cancer therapy: molecular mechanisms of action. European journal of pharmacology. 2014;740:364–378.

48. Kampan NC, Madondo MT, McNally OM, Quinn M, Plebanski M. Paclitaxel and its evolving role in the management of ovarian cancer. BioMed research international. 2015;2015.

49. Liu D, Kaufmann GF, Breitmeyer JB, Dickson KA, Marsh DJ, Ford CE. The Anti-ROR1 Monoclonal Antibody Zilovertamab Inhibits the Proliferation of Ovarian and Endometrial Cancer Cells. Pharmaceutics. 2022;14(4):837.

50. Guo J, Zhao C, Yao R, Sui A, Sun L, Liu X, et al. 3D culture enhances chemoresistance of ALL Jurkat cell line by increasing DDR1 expression. Experimental and Therapeutic Medicine. 2019;17(3):1593–1600.

51. Nowacka M, Sterzynska K, Andrzejewska M, Nowicki M, Januchowski R. Drug resistance evaluation in novel 3D in vitro model. Biomedicine & Pharmacotherapy. 2021;138:111536.

52. Anvari S, Nambiar S, Pang J, Maftoon N. Computational models and simulations of cancer metastasis. Archives of Computational Methods in Engineering. 2021; p. 1–23.

53. Cortesi M, Giordano E. Non-destructive monitoring of 3D cell cultures: new technologies and applications. PeerJ. 2022;10:e13338.

54. Liliopoulos SG, Stavrakakis GS, Dimas KS. Advanced non-linear mathematical model for the prediction of the activity of a putative anticancer agent in human-to-mouse cancer xenografts. Anticancer Research. 2020;40(9):5181–5189.

55. Jarrett AM, Hormuth II DA, Wu C, Kazerouni AS, Ekrut DA, Virostko J, et al. Evaluating patient-specific neoadjuvant regimens for breast cancer via a mathematical model constrained by quantitative magnetic resonance imaging data. Neoplasia. 2020;22(12):820–830.

56. Emert BL, Cote CJ, Torre EA, Dardani IP, Jiang CL, Jain N, et al. Variability within rare cell states enables multiple paths toward drug resistance. Nature biotechnology. 2021;39(7):865–876.

