## Supplementary figures and images for "Model Parameter identification using 2D vs 3D experimental data: a comparative analysis"

### Supplementary Figure 1

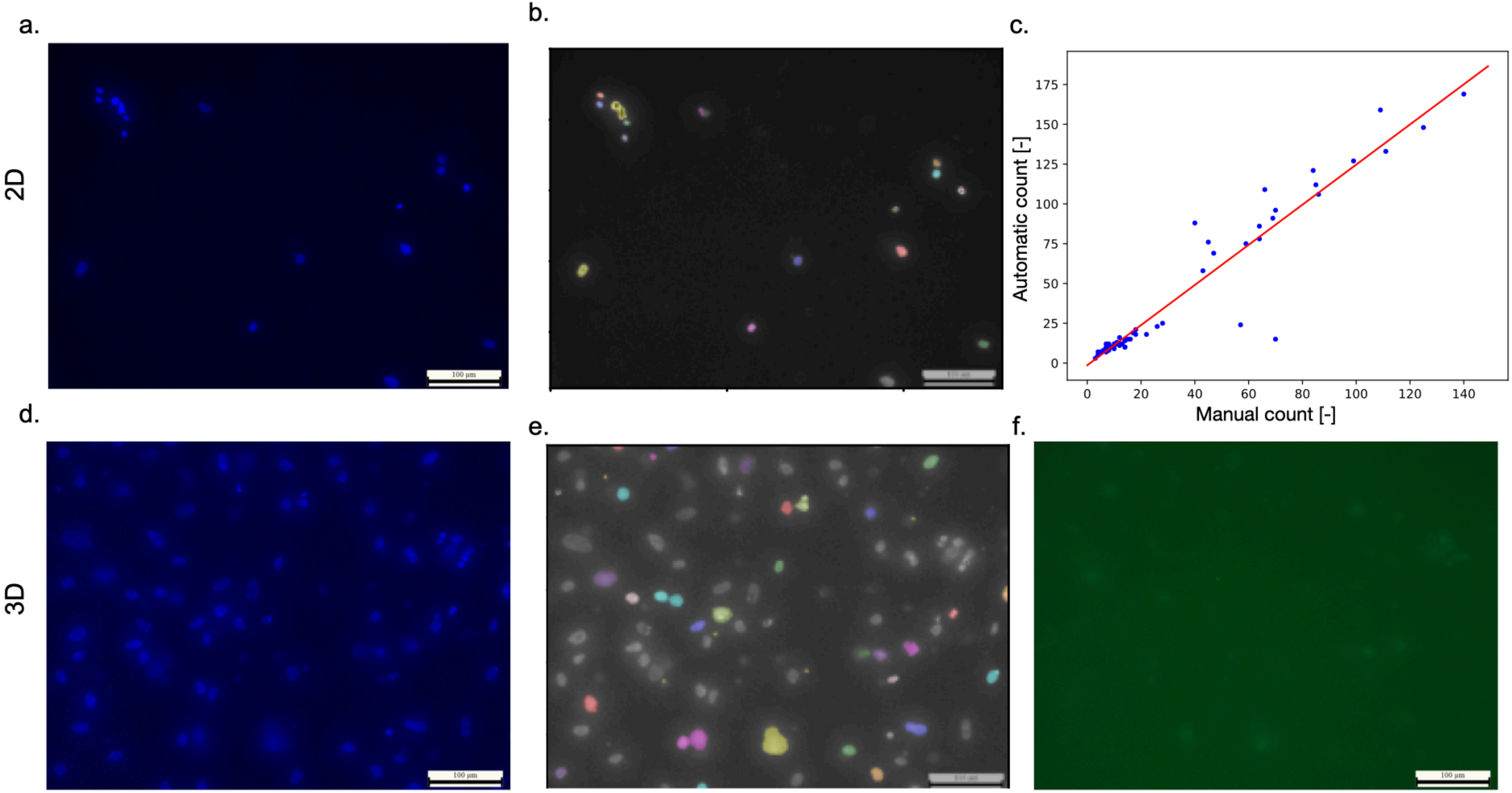

### Supplementary Figure 2

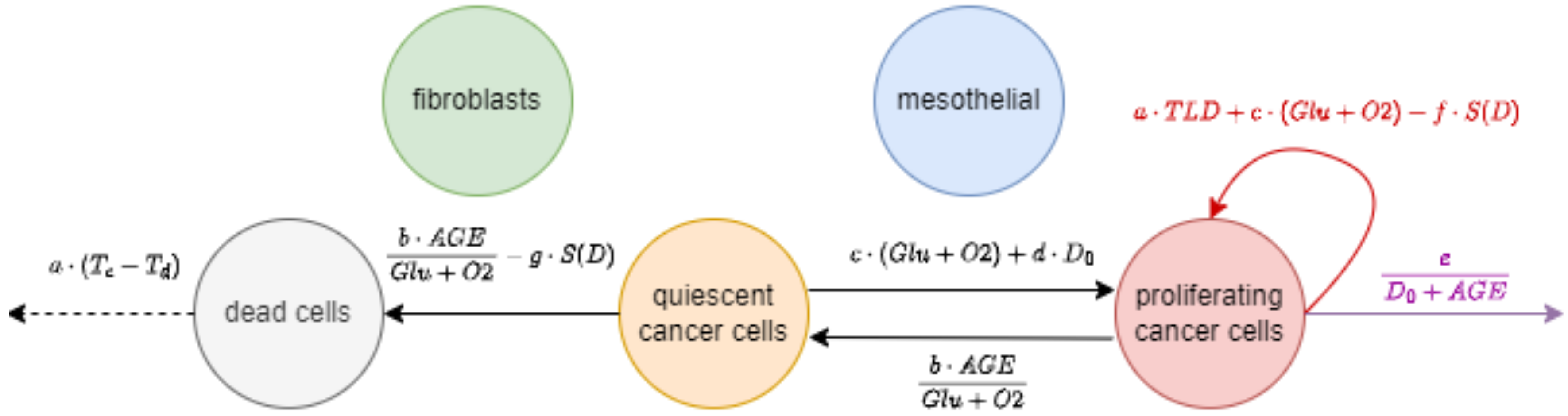

### Supplementary Figure 3

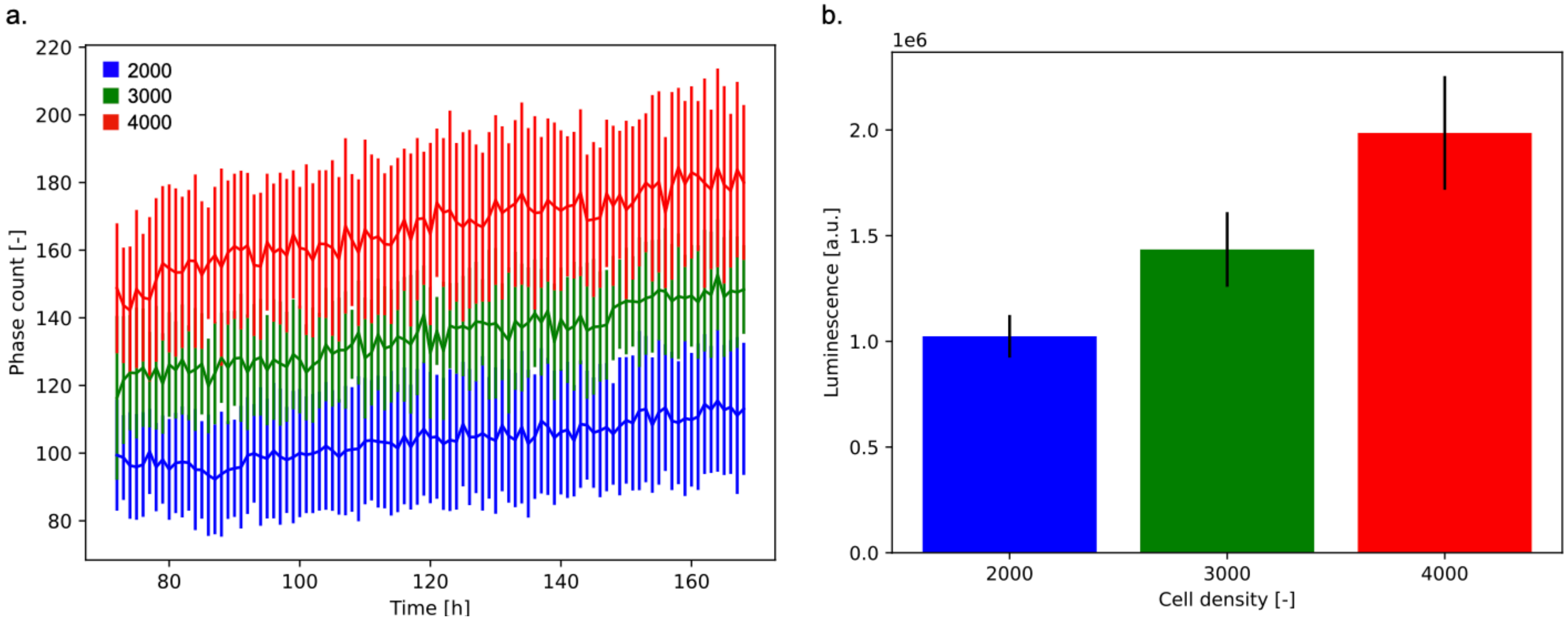
