## Supplementary data (SALSA configuration file) for "Model Parameter identification using 2D vs 3D experimental data: a comparative analysis"

### *Supplementary Material*

#### 1. Supplementary Data

##### 1.1. SALSA configuration file

In the following the configuration file used to describe the organotypic model in SALSA is reported.

```
MODEL:

 Cell Types:

 Cell Line = organoid
 1 = Dead
 2= Cancer_Quiescent
 3 = Cancer_Replication
 4 = Fibroblasts
5 = Mesothelial

Rules:

0 = 1->0, a*(TIME-TD)
1 = 1->1, 1-(a*(TIME-TD))
2 = 1->0, environment (Glc)
3 = 1->0, environment (O2)
4 = 2-> 1, b*(AGE)/(Glc+O2) - f*DRUG1
5 = 2-> 3, c*(Glc+O2) + d*D0
6 = 2-> 2, 1-(b*(AGE)/(Glc+O2)-f*DRUG1) - (c*(Glc+O2)+d*D0)
7 = 2->0.5*U, environment (Glc)
8 = 2->0.5*U, environment (O2)
9 = 3->2, b*(AGE)/(Glc+O2)
10 = 3->3+3, a*TLD+ c*(O2+Glc) -g*DRUG1
11 = 3->0+3, e/(D0+AGE)
12 = 3->3, 1-(b*(AGE)/(Glc+O2))-(a*TLD+c*(Glc+O2)-g*DRUG1)-(e/(D0+AGE))
13 = 3->U, environment (Glc)
14 = 3->U, environment (O2)
15 = 4 ->4, 1
16 = 4->0.5*U, environment (Glc)
17 = 4->0.5*U, environment (O2)
18 = 5 ->5, 1
19 = 5->0.5*U, environment (Glc)
20 = 5->0.5*U, environment (O2)

Scaffold:

material = organoid
side [cm] = 0.6
layers = 10
porosity [%] = 87

INITIAL CONDITIONS:
```

Cell Types:

1 = 0
2 = 14
3 = 0
4 = 14
5 = 71
total=28K

Scaffold:

seeding = Organoid
iterations = 72
media = RPMI
media replace frequency [day]= 3
volume per scaffold [ml] = 0.2
flow rate [ml/h] = 0

Treatment:

ID = 1
name = **drug_name**
dose [ug/ml] = **drug_dose**
treatment = [0]
type = media
